## Supplementary Information for "FBXO24 deletion causes abnormal accumulation of membraneless electron-dense granules in sperm flagella and male infertility"

[1]Research Institute for Microbial Diseases, Osaka University, 3-1 Yamadaoka, Suita, Osaka 5650871 JAPAN.
[2]Graduate School of Pharmaceutical Sciences, Osaka University, 1-6 Yamadaoka, Suita, Osaka 5650871 JAPAN.
[3]The Institute of Medical Science, The University of Tokyo, 4-6-1 Shirokanedai, Minato-ku, Tokyo 1088639 JAPAN.
[4]Center for Infectious Disease Education and Research (CiDER), Osaka University, 2-8 Yamadaoka, Suita, Osaka 5650871 JAPAN.
[5]Center for Advanced Modalities and DDS (CAMaD), Osaka University, 2-8 Yamadaoka, Suita, Osaka 5650871 JAPAN.
*Corresponding authors: Haruhiko Miyata and Masahito Ikawa

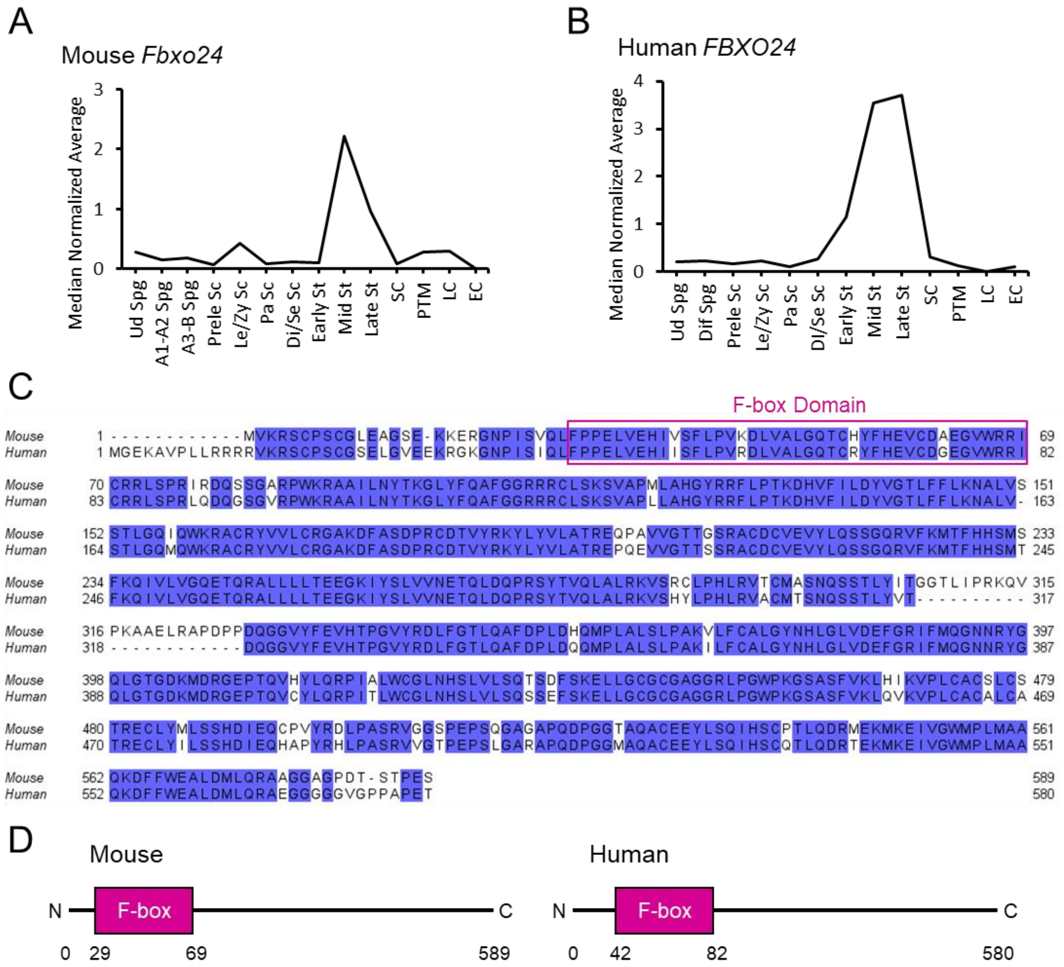


**Figure S1. Characterization of mouse and human FBXO24.**
(*A*) Expression profile of mouse *Fbxo24* in spermatogenic cells is shown. Expression profiling was obtained from the testis single cell RNA-seq data set (Hermann et al., 2018). Ud spg: Undifferentiated spermatogonia, A1- A2 Spg: A1 and A2 differentiating spermatogonia, A3-B Spg: A3, A4, In, and B differentiating spermatogonia, Prele Sc: preleptotene spermatocytes, Le/Zy Sc: leptotene/zygotene spermatocytes, Pa Sc: pachytene spermatocytes, Di/Se Sc: diplotene/secondary spermatocytes, Early St: early round spermatids, Mid St: mid round spermatids, Late St: late round spermatids, SC: Sertoli cells, PTM: peritubular myoid cells, LC: Leydig cells, and EC: Endothelial cells. (*B*) Expression of human *FBXO24* in spermatogenic cells is shown. Expression profiling was obtained from the testis single cell RNA-seq data set (Hermann et al., 2018). Dif Spg: differentiating spermatogonia. (*C*) Pairwise sequence alignment of amino acids between mouse and human FBXO24. Dark blue highlight indicates identical amino acids. (*D*) The F-box domain of mouse and human FBXO24 was identified with a protein database search (SMART; http://smart.embl-heidelberg.de/).

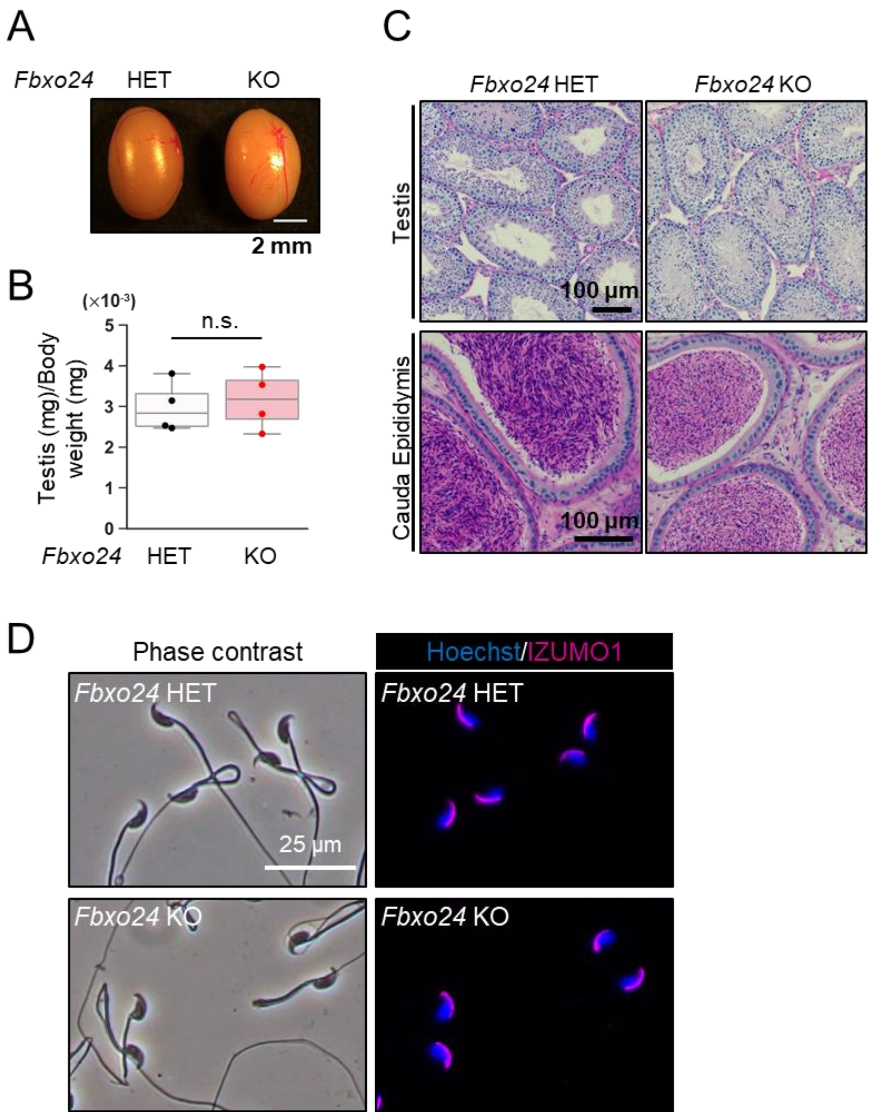


**Figure S2. Histological analyses of *Fbxo24* KO testis and epididymis.**
(*A*) Gross morphology of testes obtained from *Fbxo24* heterozygous and KO males. (*B*) Test weight (mg)/body weight (mg) of *Fbxo24* heterozygous and KO mice. (*C*) PAS-hematoxylin staining of testes and cauda epididymis sections. (*D*) Spermatozoa obtained from the cauda epididymis were stained for IZUMO1 (magenta). Nuclei were stained with Hoechst 33342 (blue).

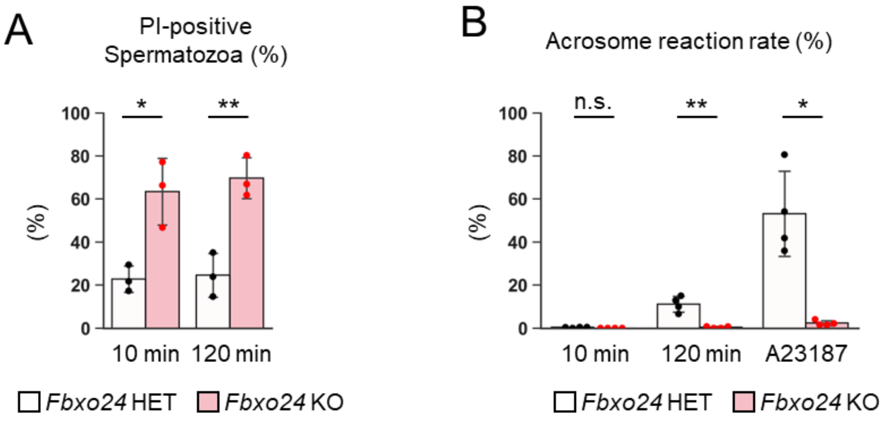


C

Quantitative value (Normalized total spectra)

| Protein | WT #1 | WT #2 | WT #3 | WT average | KO #1 | KO #2 | KO #3 | KO average |
|---|---|---|---|---|---|---|---|---|
| SNAP23 | 2 | 3 | 1 | 2 ± 1.0 | 2 | - | 2 | 2 ± 0.1 |
| STX2 | 2 | - | - | 2 | 2 | - | - | 2 |
| STX12 | 7 | 3 | 2 | 4 ± 2.6 | 4 | 2 | - | 3 ± 1.5 |
| VAMP3 | 3 | 2 | 2 | 2 ± 0.5 | 2 | 3 | 3 | 3 ± 0.5 |
| PLCD4 | 1 | 3 | 2 | 2 ± 1.1 | - | 1 | 1 | 1 ± 0 |
| IZUMO1 | 29 | 20 | 20 | 23 ± 5.2 | 38 | 11 | 11 | 20 ± 15.6 |

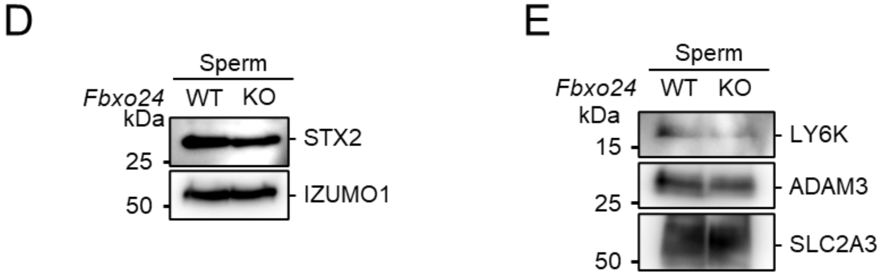


**Figure S3. Lack of *Fbxo24* impairs sperm viability and the acrosome reaction.** (*A*) Viability of cauda epididymal spermatozoa was assessed with propidium iodide (PI) at 10 min and 120 min incubation in TYH medium. (*B*) The acrosome reaction rates of cauda epididymal spermatozoa were assessed using RBGS Tg mice after 10 min and 120 min of incubation in TYH medium. To induce the acrosome reaction, $Ca^{2+}$ ionophore A23187 was added to the medium after 120 min of incubation. (*C*) MS analyses of triton X-100 soluble proteins obtained from mature spermatozoa. Quantitative value of identified SNARE-related proteins and PLCD4 were listed. (*D*) Immunodetection of STX2 in *Fbxo24* WT and KO spermatozoa. IZUMO1 was used as loading control. (*E*) Immunodetection of LY6K and ADAM3 in *Fbxo24* WT and KO spermatozoa. SLC2A3 was used as loading control.

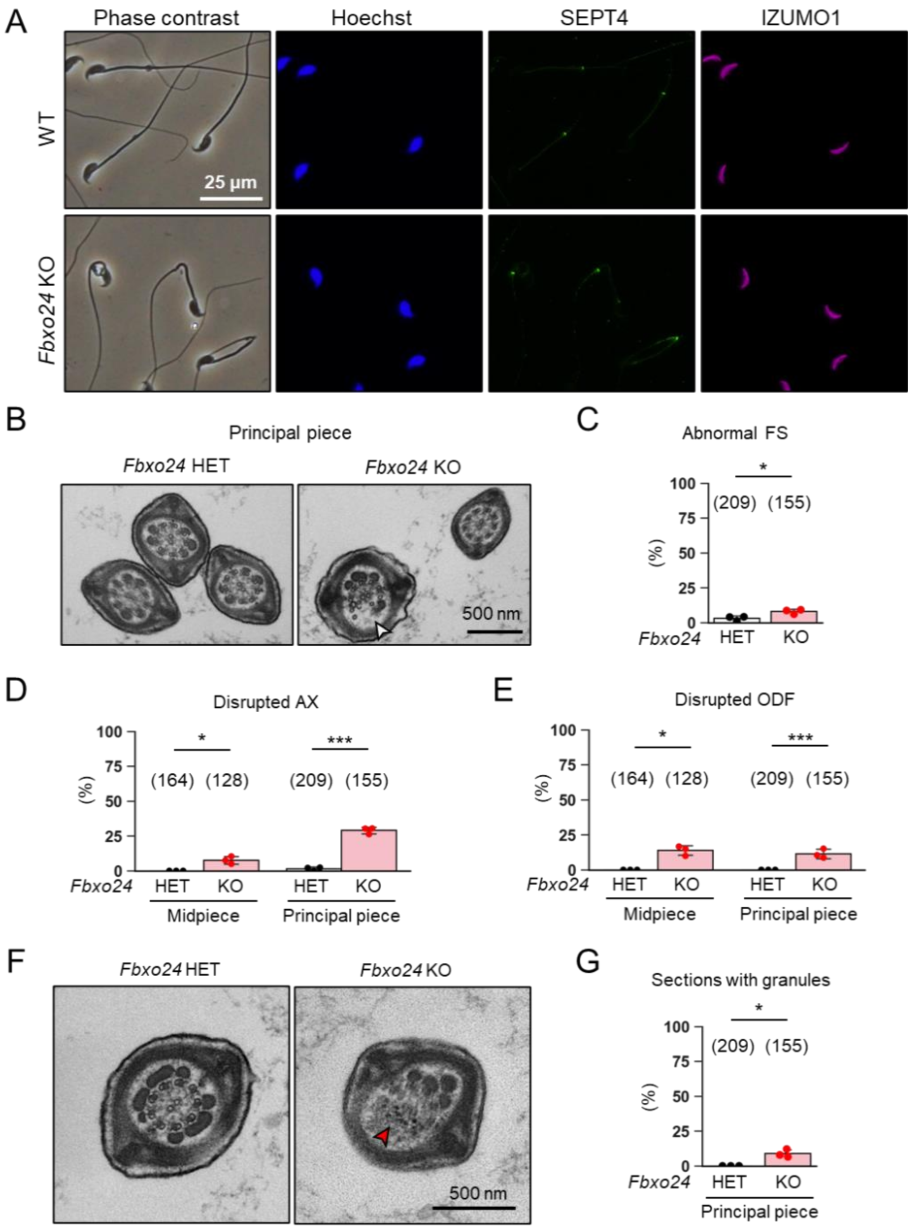


**Figure S4. Sperm flagellum ultrastructure was disorganized in *Fbxo24* KO mice.**
(*A*) Localization of the annulus in cauda epididymal spermatozoa. Anti-SEPT4 antibody was used to visualize the annulus (green). Nuclei were detected with Hoechst 33342 (blue). The acrosome was detected with anti-IZUMO1 antibody (magenta). (*B*) Transmission electron microscopy (TEM) observation of the principal piece. A white arrowhead indicates a disruption of the axoneme and outer dense fiber (ODF). (*C*) Percentages of morphologically abnormal fibrous sheaths (FS) observed with TEM. The number of flagellar sections analyzed is shown above each bar. (*D*) Percentages of morphologically abnormal axonemes (AX) observed with TEM. The number of flagellar sections analyzed are shown above each bar. (*E*) Percentages of morphologically abnormal ODF observed with TEM. The number of flagellar sections analyzed are shown above each bar. (*F*) TEM observation of the principal piece. A red arrowhead indicates electron-dense granules. (*G*) Percentages of electron-dense granules observed in principal piece cross sections. The number of flagellar sections analyzed are shown above each bar.

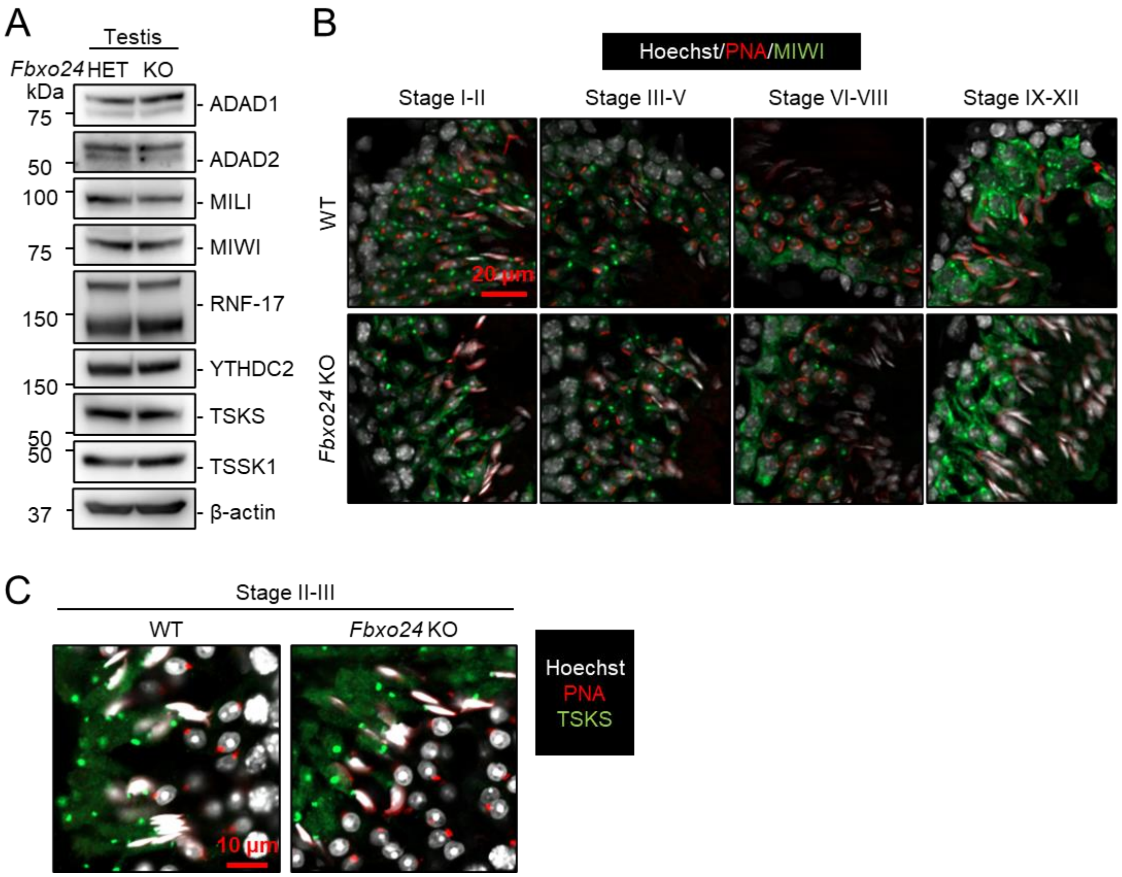


**Figure S5. No clear differences were found in the amount and localization of RNP granule-related proteins between control and *Fbxo24* KO testes.**
(*A*) Immunodetection of ADAD1, ADAD2, MILI, MIWI, RNF-17, YTHDC2, TSKS and TSSK1 in *Fbxo24* heterozygous and KO testes. β-actin was used as loading control. (*B*) Immunofluorescence observation of MIWI (green) in WT and *Fbxo24* KO testes. Acrosomes were stained with PNA (red) and nuclei were stained with Hoechst 33342 (white). (*C*) Immunofluorescence observation of TSKS (green) in WT and *Fbxo24* KO testes. Acrosomes were stained with PNA (red) and nuclei were stained with Hoechst 33342 (white).

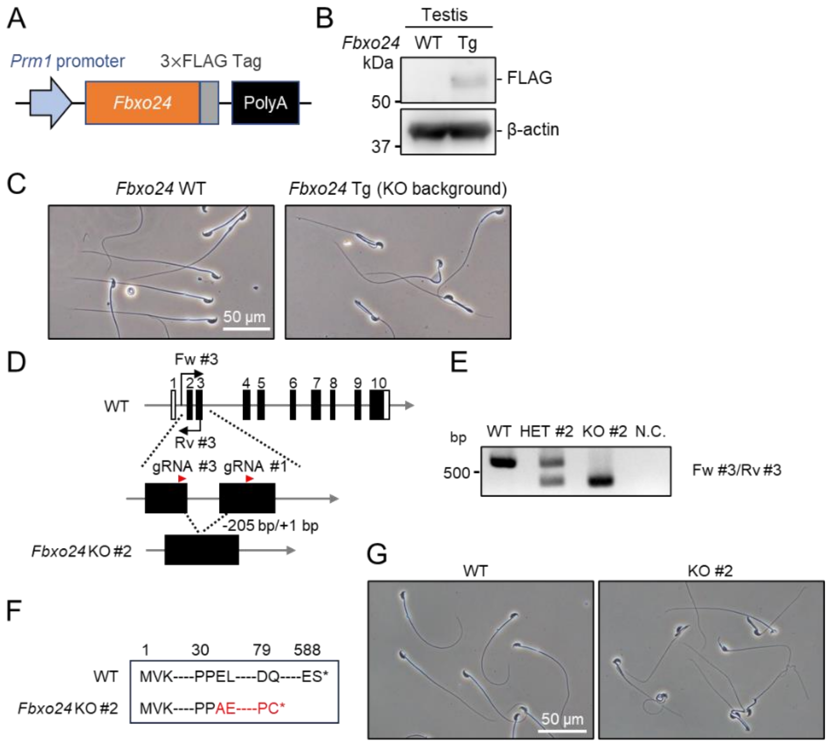


H

| Identified Proteins | Quantitative Value (Normalized Total Spectra) | |
|---|---|---|
| | Tg | WT |
| FBXO24 | 94 | 0 |
| IPO5 | 68 | 0 |
| STIP1 | 54 | 0 |
| HSPA1B | 41 | 0 |
| HSPA4 | 36 | 0 |
| SUGT1 | 34 | 0 |
| DNAJC7 | 28 | 0 |
| FKBP4 | 27 | 0 |
| TUBB3 | 25 | 0 |
| RPAP3 | 25 | 0 |
| RUVBL2 | 24 | 0 |
| SKP1 | 21 | 0 |
| RUBCL1 | 21 | 0 |
| DNAJA4 | 20 | 0 |
| NUDCD3 | 20 | 0 |

**Figure S6. Generation and analyses of *Fbxo24-3×FLAG* Tg mice.**
(*A*) Schematic of the *Fbxo24-3×FLAG* construct used to generate Tg mice. (*B*) Immunodetection of FBXO24-3×FLAG in the testis from the Tg mice. β-actin was used as a loading control. (*C*) Morphology of cauda epididymal spermatozoa. (*D*) Schematic for generating *Fbxo24* KO #2 mice using the CRISPR/Cas9 system. White boxes indicate untranslated regions while black boxes indicate protein coding regions. The gRNAs used are shown. Fw and Rv indicate the forward and reverse primer used for genotyping, respectively. (*E*) Genotyping of obtained *Fbxo24* mutant mice. Fw #3-Rv #3 primers in Fig. S6D were used. N.C. indicates negative control (water). *(F)* Predicted FBXO24 amino acid sequences of KO #2 mutant mice. The 205 bp deletion and 1 bp insertion resulted in E32D mutation with a premature stop codon introduced 49 amino acids later. (G) Morphology of mature spermatozoa obtained from cauda epididymis. (*H*) The list of identified proteins by IP-MS analysis. The top 15 identified proteins in Tg are shown. The Quantitative Value (Normalized Total Spectra) was calculated using Scaffold proteome software. IgG-related proteins, which is likely from the antibody used for IP, were removed from the list.

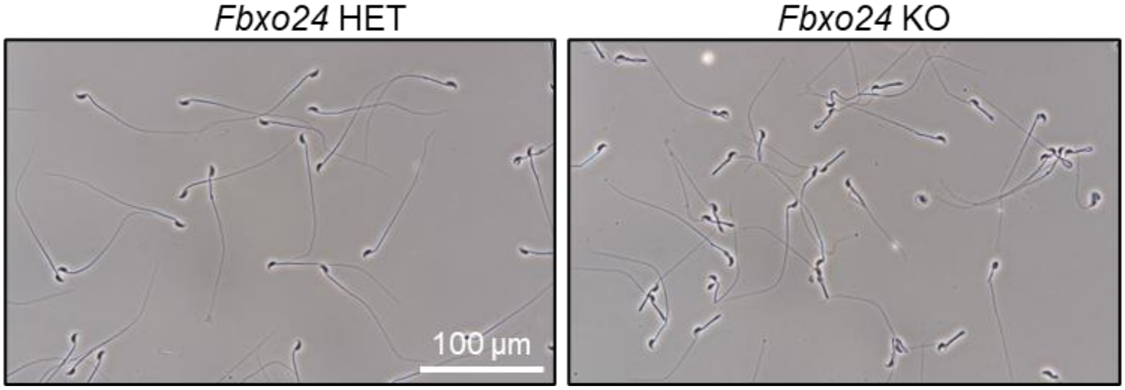


**Figure S7. Spermatogenic cells obtained from cauda epididymis.**
Spermatozoa were obtained *Fbxo24* heterozygous or KO male mice. Immature spermatogenic cells or somatic cells were rarely observed.

**Table S1. Primers and gRNAs used in this study.**

| primer | sequence (5'-3') |
|---|---|
| *Fbxo24 Fw #1* | tcctctggcaggtgaagcgc |
| *Fbxo24 Fw #2* | gatgtgttgattgtatgagacttcc |
| *Fbxo24 Fw #3* | actagcagcaagaagtccagg |
| *Fbxo24 Rv #1* | tctcgctcatgatagtcagg |
| Fbxo24 Rv #2 | gaagtatggcaaagtcactgagtgc |
| Fbxo24 Rv #3 | aagaatggcagctctcttcc |
| *Fbxo24 Tg Fw* | tgaagtgtgtgatgctgagg |
| *Fbxo24 Tg Rv* | gttcccgagtagccaaaaca |
| *Fbxo24 Cloning Fw* | atggatccgccgccatggtgaagcgcagctgcccttc |
| *Fbxo24 Cloning Rv* | gcgaattcgctctcgggagtcgaggtgtctgg |
| *Skp1 Cloning Fw* | attctagagccgccatgcctacgataaagttgc |
| *Skp1 Cloning Rv* | gcctgcagcttctcttcacaccattgg |
| *Ipo5 Cloning Fw* | atgtcgacgccgccatggcggcggccgcggcggagcagc |
| *Ipo5 Cloning Rv* | gcgctagcggcagagttcaggagctcctggatgg |
| *Actb Fw* | catccgtaaagacctctatgccaac |
| *Actb Rv* | atggagccaccgatccaca |

| gRNA | target sequence (5'-3') |
|---|---|
| *Fbxo24* gRNA #1 | tgtggaggcgcatctgtcga |
| *Fbxo24 gRNA #2* | tcctgaaggaagtcgagccg |
| *Fbxo24 gRNA #3* | tcagttgttccccccagagc |

**Table S2. Antibodies used in this study**

| Antibodies | Source | Identifier |
|---|---|---|
| Rabbit anti-KPNA2 | Abcam | ab84440 |
| Rabbit anti-β-actin | MBL | PM053 |
| Rabbit anti-human septin4 | IBL | 18987 |
| Rabbit anti-IPO5 (H-300) | SantaCruz | sc-11369 |
| Rabbit anti-Miwi (G82) | Cell Sinaling Technology | 2079 |
| Rabbit anti-Syntaxin2 | Synaptic Systems | 110 123 |
| Rabbit anti-LY6K | (Yoshitake et al., 2008) | |
| Rabbit anti-HA tag | MBL | 561 |
| Rabbit anti-ADAD1 | (Lu et al., 2023) | |
| Rabbit anti-TSKS | (Shimada et al., 2023) | |
| Rabbit anti-TSSK1 | (Shimada et al., 2023) | |
| Rat anti-ADAD2 | (Lu et al., 2023) | |
| Rat anti-IZUMO1 | (Ikawa et al., 2011) | |
| Rat anti-SLC2A3 | (Fujihara et al., 2012) | |
| Rat anti-PA tag NZ-1 | FUJIFILM Wako | 012-25863 |
| Mouse anti-1D4 tag | Gift from Dr. Martin M. Matzuk | |
| Mouse anti-FLAG tag (M2) | Sigma-Aldrich | F1804 |
| Mouse anti-α-tubulin (B-5-1-2) | Sigma-Aldrich | T5168 |
| Mouse anti-acetylated tubulin | Sigma-Aldrich | T7451 |
| Mouse anti-ADAM3 (F-4) | SantaCruz | sc-365288 |
| Mouse anti-IPO5 (B-7) | SantaCruz | sc-55527 |
| Mouse anti-β-actin (AC-15) | Abcam | ab6276 |
| Mouse anti-KPNB1 | Abcam | ab2811 |
| Mouse anti-phosphorylated-tyrosine | Merck Millipore | 05-321 |
| Goat anti-rabbit IgG-HRP | Jackson ImmunoResearch | 111-036-045 |
| Goat anti-rat IgG-HRP | Jackson ImmunoResearch | 112-035-167 |
| Goat anti-mouse IgG-HRP | Jackson ImmunoResearch | 115-036-062 |
| Goat anti-rabbit IgG-Alexa Fluor 488 | Thermo Fisher Scientific | A11070 |
| Goat anti-rabbit IgG-Alexa Fluor 546 | Thermo Fisher Scientific | A11071 |
| Goat anti-rat IgG-Alexa Fluor 488 | Thermo Fisher Scientific | A11006 |
| Goat anti-rat IgG-Alexa Fluor 546 | Thermo Fisher Scientific | A11081 |
| Goat anti-mouse IgG-Alexa Fluor 546 | Thermo Fisher Scientific | A11017 |
| Goat anti-mouse IgG-Alexa Fluor 546 | Thermo Fisher Scientific | A11018 |

### Legends for Movies S1 and S2

**Movie S1 (separate file).** Sperm motion of *Fbxo24* heterozygous mice was recorded after 120 min of incubation in TYH medium.

**Movie S2 (separate file).** Sperm motion of *Fbxo24* KO mice was recorded after 120 min of incubation in TYH medium.
